# A human coronavirus evolves antigenically to escape antibody immunity

**DOI:** 10.1101/2020.12.17.423313

**Authors:** Rachel Eguia, Katharine H. D. Crawford, Terry Stevens-Ayers, Laurel Kelnhofer-Millevolte, Alexander L. Greninger, Janet A. Englund, Michael J. Boeckh, Jesse D. Bloom

## Abstract

There is intense interest in antibody immunity to coronaviruses. However, it is unknown if coronaviruses evolve to escape such immunity, and if so, how rapidly. Here we address this question by characterizing the historical evolution of human coronavirus 229E. We identify human sera from the 1980s and 1990s that have neutralizing titers against contemporaneous 229E that are comparable to the anti-SARS-CoV-2 titers induced by SARS-CoV-2 infection or vaccination. We test these sera against 229E strains isolated after sera collection, and find that neutralizing titers are lower against these “future” viruses. In some cases, sera that neutralize contemporaneous 229E viral strains with titers >1:100 do not detectably neutralize strains isolated 8–17 years later. The decreased neutralization of “future” viruses is due to antigenic evolution of the viral spike, especially in the receptor-binding domain. If these results extrapolate to other coronaviruses, then it may be advisable to periodically update SARS-CoV-2 vaccines.

## Introduction

The SARS-CoV-2 pandemic has caused an urgent need to determine how well antibody immunity protects against SARS-CoV-2 infection. The evidence so far is promising. Neutralizing and anti-spike antibodies elicited by natural infection correlate with reduced SARS-CoV-2 infection of humans (Addetia et al., 2020; Lumley et al., 2020), and vaccines that elicit such antibodies protect humans with high efficacy (Polack et al., 2020). These findings in humans are corroborated by a multitude of animal studies showing that neutralizing antibodies to the SARS-CoV-2 spike protect against infection and disease (Alsoussi et al., 2020; McMahan et al., 2020; Walls et al., 2020; Zost et al., 2020).

However, humans are repeatedly re-infected with the “common-cold” coronaviruses 229E, OC43, HKU1, and NL63 (Edridge et al., 2020; Hendley et al., 1972; Schmidt et al., 1986). For instance, serological studies suggest that the typical person is infected with 229E every 2–3 years (Edridge et al., 2020; Schmidt et al., 1986), although a lower infection rate and no 229E re-infections were reported in a 4-year study that identified infections by the criteria of a positive PCR test in the context of respiratory illness (Aldridge et al., 2020). In any case, the fact that common-cold coronavirus re-infections occur at some appreciable rate has led to concerns that coronavirus immunity is not “durable.” These concerns initially focused on the possibility that the immune response itself is not durable (Ibarrondo et al., 2020). This possibility now seems less likely, as SARS-CoV-2 infection induces neutralizing antibodies and memory B cells with dynamics similar to other respiratory viruses (Crawford et al., 2020a; Gaebler et al., 2020; Rodda et al., 2020; Wajnberg et al., 2020).

But there is another mechanism by which viruses can re-infect even in the face of long-lived and effective antibodies: antigenic evolution. For example, infection with influenza virus elicits antibodies that generally protect humans against that same viral strain for at least several decades (Couch and Kasel, 1983; Yu et al., 2008). Unfortunately, influenza virus undergoes rapid antigenic evolution to escape these antibodies (Bedford et al., 2014), meaning that although immunity to the original viral strain lasts for decades, humans are susceptible to infection by its descendants within about 5 years (Couch and Kasel, 1983; Ranjeva et al., 2019). This continual antigenic evolution is the reason that the influenza vaccine is periodically updated.

Strangely, the possibility of antigenic evolution by coronaviruses has received only modest attention, perhaps because coronaviruses have lower mutation rates than other RNA viruses (Denison et al., 2011; Sanjuán et al., 2010). However, mutation rate is just one factor that shapes antigenic evolution; influenza and measles virus both have high mutation rates, but only the former undergoes rapid antigenic evolution. Furthermore, the assumption of minimal coronavirus antigenic evolution is not supported by the limited evidence to date. In the 1980s, human-challenge studies found that individuals infected with one strain of 229E were resistant to re-infection with that same strain, but partially susceptible to a different strain (Reed, 1984). Additional experimental studies suggest that sera or antibodies can differentially recognize spike proteins from different 229E strains (Shirato et al., 2012; Wong et al., 2017). From a computational perspective, several studies have reported that the spikes of 229E and OC43 evolve under positive selection (Chibo and Birch, 2006; Kistler and Bedford, 2020; Ren et al., 2015), which is often a signature of antigenic evolution.

Here we experimentally assess whether coronavirus 229E escapes neutralization by human polyclonal sera by reconstructing the virus’s evolution over the last several decades. We show that historical sera that potently neutralize virions pseudotyped with contemporary 229E spikes often have little or no activity against spikes from 229E strains isolated 8–17 years later. Conversely, modern sera from adults generally neutralize spikes from a wide span of historical viruses, whereas modern sera from children best neutralize spikes from recent viruses that circulated during the children’s lifetimes. These patterns are explained by antigenic evolution of the spike, especially within the receptor-binding domain. If SARS-CoV-2 undergoes similarly rapid antigenic evolution, then it may be advisable to periodically update vaccines to keep pace with viral evolution.

## Results

### Phylogenetic analysis of 229E spikes to identify historical strains for experimental study

We focused our studies on the viral spike protein because it is the main target of neutralizing antibodies (Tortorici and Veesler, 2019), and because anti-spike antibodies are the immune parameter best established to associate with protection against coronavirus infection in humans (Addetia et al., 2020; Callow, 1985; Callow et al., 1990; Lumley et al., 2020; Polack et al., 2020).

Because SARS-CoV-2 has circulated in humans for less than a year, we needed to choose another coronavirus with a more extensive evolutionary history. Of the four human-endemic common-cold coronaviruses, the two alphacoronaviruses 229E and NL63 are similar to SARS-CoV-2 in binding a protein receptor via the spike’s receptor-binding domain (RBD, also known as S1-B) (Hofmann et al., 2005; Walls et al., 2020; Yeager et al., 1992). In contrast, the two betacoronaviruses OC43 and HKU1 bind glycan receptors via the spike’s N-terminal domain (NTD, also known as S1-A) (Hulswit et al., 2019). Antibodies that block receptor binding dominate the neutralizing activity of immunity elicited by SARS-CoV-2 infection (Abe et al.; Piccoli et al., 2020; Tan et al., 2020), so we reasoned that even though SARS-CoV-2 is a betacoronavirus, its antigenic evolution is more likely to be foreshadowed by the two human alphacoronaviruses that also use their RBD to bind a protein receptor. Of these two viruses, we chose 229E since it has circulated in humans for >50 years (Hamre and Procknow, 1966), whereas NL63 was only identified in 2003 (van der Hoek et al., 2004).

We inferred a phylogenetic tree of 229E spikes from direct or low-passage human isolates (Figure 1A), excluding older strains passaged extensively in the lab (Hamre and Procknow, 1966). There are several important features of the tree. First, it is clock-like, with sequence divergence proportional to virus isolation date (Figures 1A and S1). Second, the tree is “ladder-like,” with short branches off a single trunk (Figure 1A). The ladder-like shape of the 229E phylogeny has been noted previously (Chibo and Birch, 2006; Hodcroft, 2020; Kistler and Bedford, 2020), and is a signature of viruses such as influenza for which immune pressure drives population turnover by selecting for antigenic variants (Bedford et al., 2011; Fitch et al., 1997). Third, sequences group by date rather than country of isolation (in Figure 1A, sequences from different countries but the same year are nearby). Phylogenies that organize by date rather than geography indicate fast global transmission, another signature of human influenza virus (Lemey et al., 2014; Nelson et al., 2007). Finally, although there is some intra-spike recombination, it is among closely related strains and does not affect the broader topology of the tree (Figure S2). For our study, the key implication of the above observations is that date of virus isolation is a good proxy for evolutionary position, since 229E evolves primarily along a single trajectory through time.

**Figure 1.**
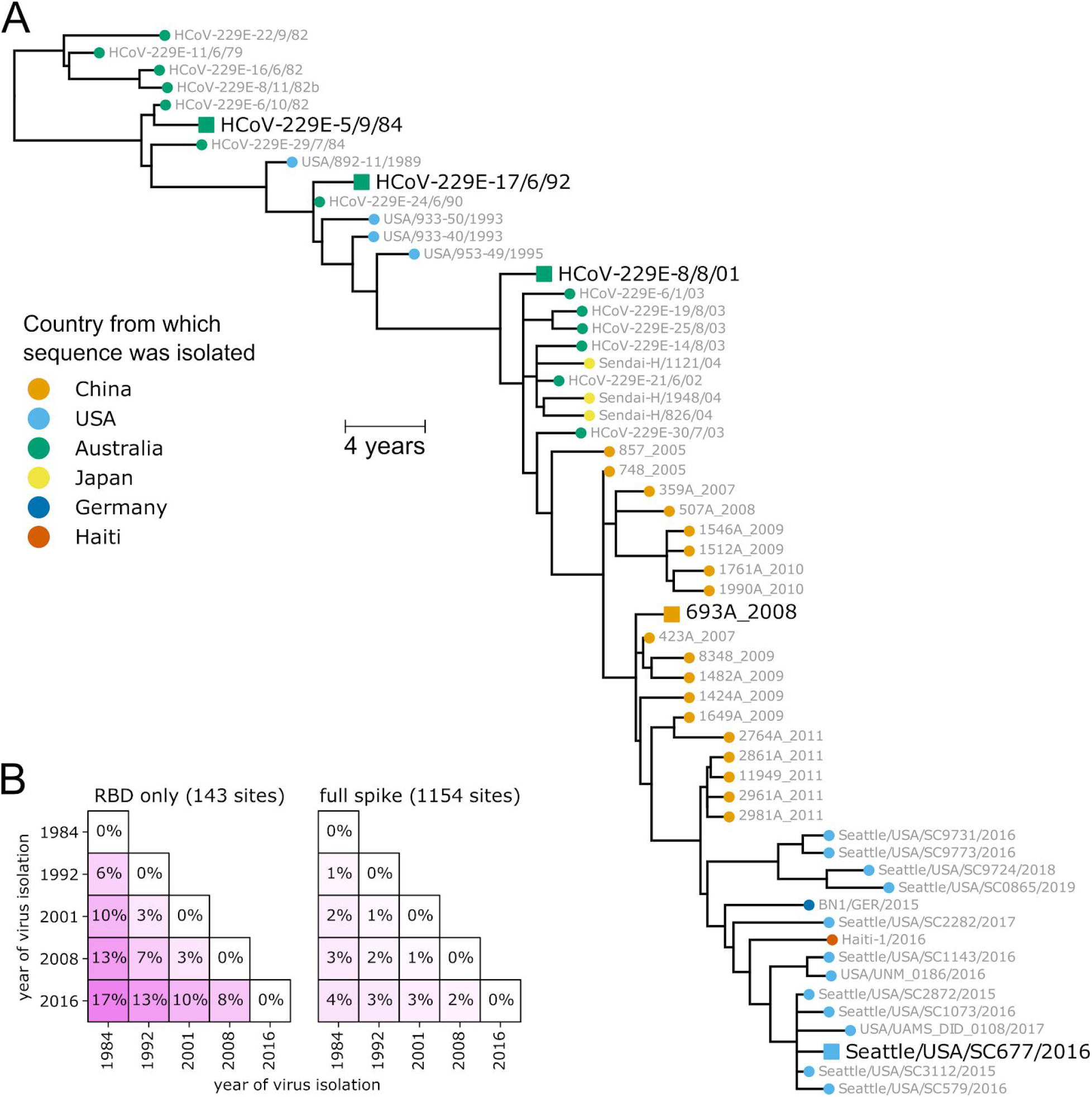
Spikes used in this study. (A) Phylogenetic tree of 229E spikes, with tips colored by the country from which the virus was isolated. The spikes used in the experiments are indicated with black text and square shapes. The tree is a maximum-likelihood inference with IQ-TREE (Minh et al., 2020) with a codon-substitution model and re-scaled with TreeTime (Sagulenko et al., 2018) to position tips by to date of isolation. Figure S1 shows a tree with branch lengths proportional to divergence rather than time, and validates clock-like evolution. Figure S2 shows recombination does not substantially affect the phylogenetic placements of the spikes used in the experiments. (B) Protein sequence divergence of the spikes used in the experiments, computed over just the receptor-binding domain (RBD) or the full sequence. Divergence is the Levenshtein distance between the amino-acid sequences divided by the number of sites.

For our experiments, we chose five spikes from 229E viruses spaced at roughly 8-year intervals spanning 1984 to 2016 (Figure 1A). We synthesized genes encoding all five spikes, truncating the last 19 residues of the cytoplasmic tail since this improves titers of spike-pseudotyped viruses (Crawford et al., 2020a; Kawase et al., 2009; Rogers et al., 2020). These five spikes differ by up to 4% in amino-acid sequence over their entire lengths, but are vastly more different in their RBDs, with 17% RBD divergence between 1984 and 2016 (Figure 1B). We generated lentiviral particles pseudotyped with each spike, and found that all five supported high infectious titers in cells expressing 229E’s receptor aminopeptidase N (Yeager et al., 1992) and the activating protease TMPRSS2 (Bertram et al., 2013) (Figure S3). Any major antigenic evolution by 229E since the 1980s should be captured by differences among these five spikes.

### Neutralizing titers of historical sera drop rapidly against spikes from “future” viruses

To test if the 229E spikes had evolved to escape neutralization by human immunity, we used historical sera collected from adults between 1985 and 1990. Since the typical person is infected with 229E every 2–5 years (Aldridge et al., 2020; Edridge et al., 2020; Schmidt et al., 1986), many of these individuals should have been infected with 1984-like viruses within a few years preceding sera collection. None of the individuals would have been infected with any of the later viruses, since those viruses did not yet exist at the time of sera collection.

Nearly all sera collected from 1985–1990 had at least some neutralizing activity against viral particles pseudotyped with the 1984 spike (25 of 27 sera had titers >1:10; Figures S4 and S5). We focused further analysis on the roughly 30% of sera (8 of 27) that had neutralizing titers against the 1984 spike of >1:90 (Figure S5). Our reason for focusing on these sera is that their anti-229E neutralizing titers are comparable to anti-SARS-CoV-2 sera neutralizing titers several months after recovery from COVID-19 (Crawford et al., 2020a; Wajnberg et al., 2020) or receipt of the Moderna mRNA-1273 vaccine (Widge et al., 2020).

All sera that potently neutralized virions pseudotyped with the 1984 spike had reduced titers against more recent spikes (Figure 2A). In some cases, the drop in neutralization of more recent spikes was dramatic. For instance, serum collected from a 28-year old in 1990 neutralized the 1984 spike at a titer of 1:125 but did not neutralize the 1992 spike at our limit of detection of 1:10 (Figure 2A). Similarly, serum collected from a 24-year old in 1987 neutralized the 1984 spike at 1:120 but barely neutralized the 1992 spike and did not detectably neutralize spikes more recent than 1992 (Figure 2A). Only 2 of 8 sera that potently neutralized the 1984 spike detectably neutralized all subsequent spikes, and only at greatly reduced titers against the most recent spikes.

**Figure 2.**
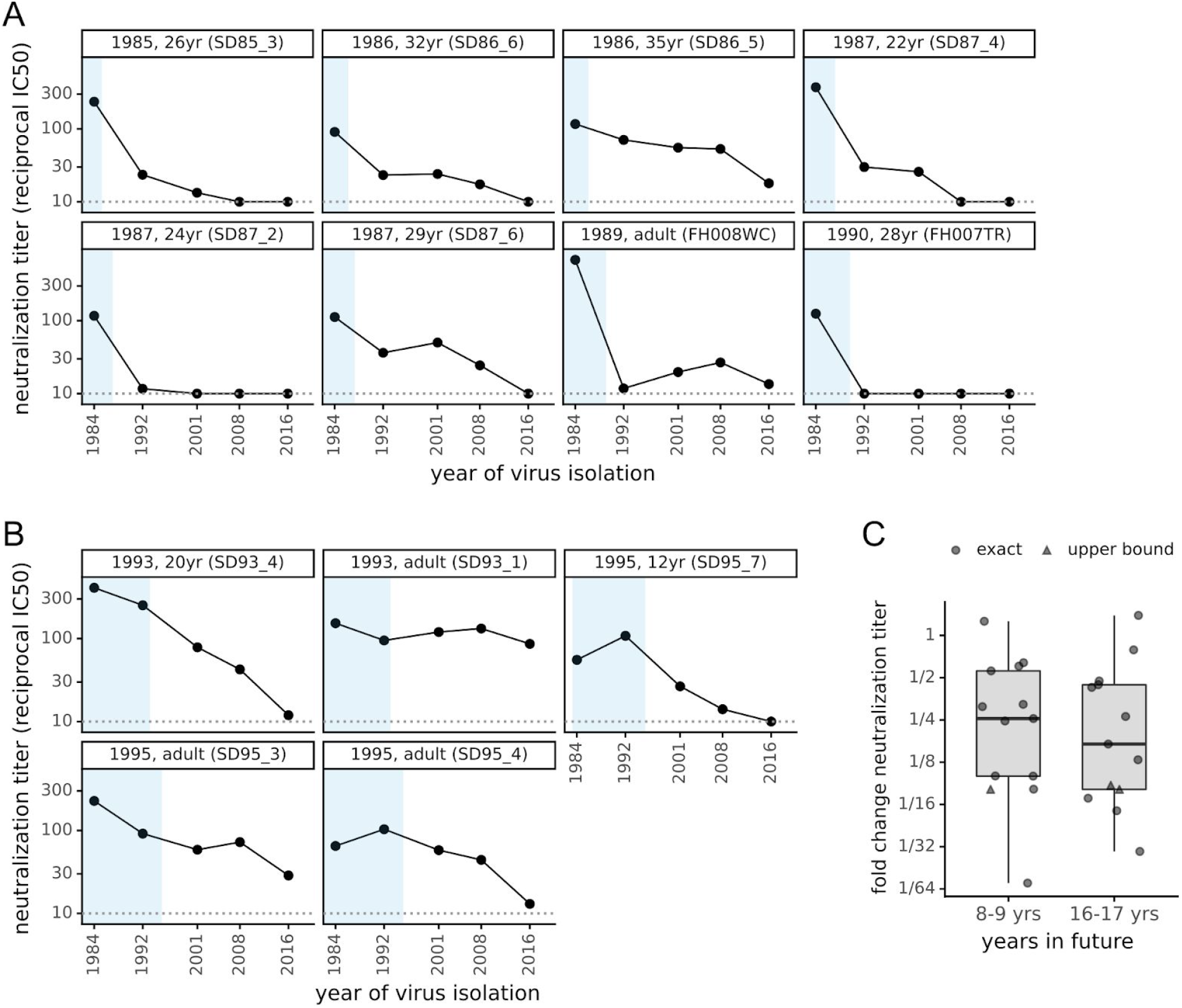
The neutralizing activity of human sera is lower against “future” viruses than those that elicited the immunity. (A) Sera collected between 1985 and 1990 was tested in neutralization assays against spikes from viruses isolated between 1984 and 2016. Each plot facet is a different serum, and black points show its neutralizing titer against viruses from the indicated year. Blue shading indicates the portion of plotted timeframe during which the individual could have been infected prior to serum collection. The dotted gray horizontal line indicates the limit of detection (titer of 1:10). Plot titles give the year of serum collection, the individual’s age when the serum was collected, and the serum ID. (B) Plots like those in (A) but for sera collected between 1993 and 1995. (C) The fold change in neutralization titer against viruses isolated 8–9 or 16–17 years in the “future” relative to the virus isolated just before the serum was collected. Box plots show the median and interquartile range, and each point is the fold change for a single serum. For a few sera (triangles), the fold change is censored (as an upper bound) because the titer against the future virus was below the limit of detection.

To confirm that these results reflect antigenic evolution rather than some unique neutralization susceptibility of the 1984 spike, we repeated similar experiments using sera collected from 1992–1995 and initially screening for neutralization of the 1992 spike. Again, nearly all (18 of 19) sera detectably neutralized the 1992 spike, with about a quarter (5 of 19) having titers >1:90 (Figures S4 and S5). These potent sera also neutralized the older 1984 spike with high titers, but again generally had lower activity against spikes from viruses isolated after the sera were collected (Figure 2B). Together, the results from the two sera collection timeframes indicate that the 229E spike is evolving antigenically, such that immunity elicited by infection with prior viruses is often ineffective at neutralizing future viruses.

To quantify the rate of antigenic evolution, for all sera in Figure 2A,B we computed the drop in neutralization titer against spikes from one and two timepoints later relative to the contemporaneous spike. The median drop in titer was 4-fold against viruses from 8–9 years in the future, and >6-fold for viruses 16–17 years in the future (Figure 2C). However, these medians mask substantial serum-to-serum variation in neutralization of antigenically evolved future viruses. Neutralization by some sera is eroded >10-fold by just 8–9 years of viral evolution, whereas neutralization by a few sera is unaffected even by 16–17 years of evolution.

### Modern sera neutralize viruses that circulated throughout an individual’s lifetime

The above results show that viral antigenic evolution erodes the capacity of anti-229E immunity to neutralize the future descendants of the viruses that elicited the immunity. We next addressed a related question: does serum immunity durably retain the capacity to neutralize historical 229E strains that an individual was exposed to many years ago?

To address this question, we used modern sera collected in 2020 from children and adults. The adults were alive during circulation of all five 229E spikes in our panel (i.e., they were born before 1984), but the children could only have been exposed to the more recent spikes. We screened 31 modern sera against the 2016 spike, and found that 25 of 31 detectably neutralized at a threshold of 1:10 (Figures S4 and S5). We again focused further analysis on the more potent sera with titers >1:90 (7 of 31 sera, Figure S5).

Modern adult sera that potently neutralized the 2016 spike also neutralized all prior spikes dating back to 1984 (Figure 3). In contrast, the children’s sera neutralized spikes from viruses that circulated during the children’s lifetimes but often had reduced activity against spikes from before the children were born (Figure 3). However, neutralization by children’s sera generally extends further “back in time” to viruses that preceded birth than neutralization by adult sera in Figure 2A,B extends “forward in time” to viruses that circulated after the sera was collected. Similar time asymmetry in antigenic evolution has been described for influenza virus (Sandbulte et al., 2011). Overall, the results in Figure 3 show that neutralizing immunity can encompass the entire spectrum of spikes an individual has been exposed to, consistent with the notion that reduced neutralization of future viruses is due to antigenic evolution rather than a lack of durability in immunity itself.

**Figure 3.**
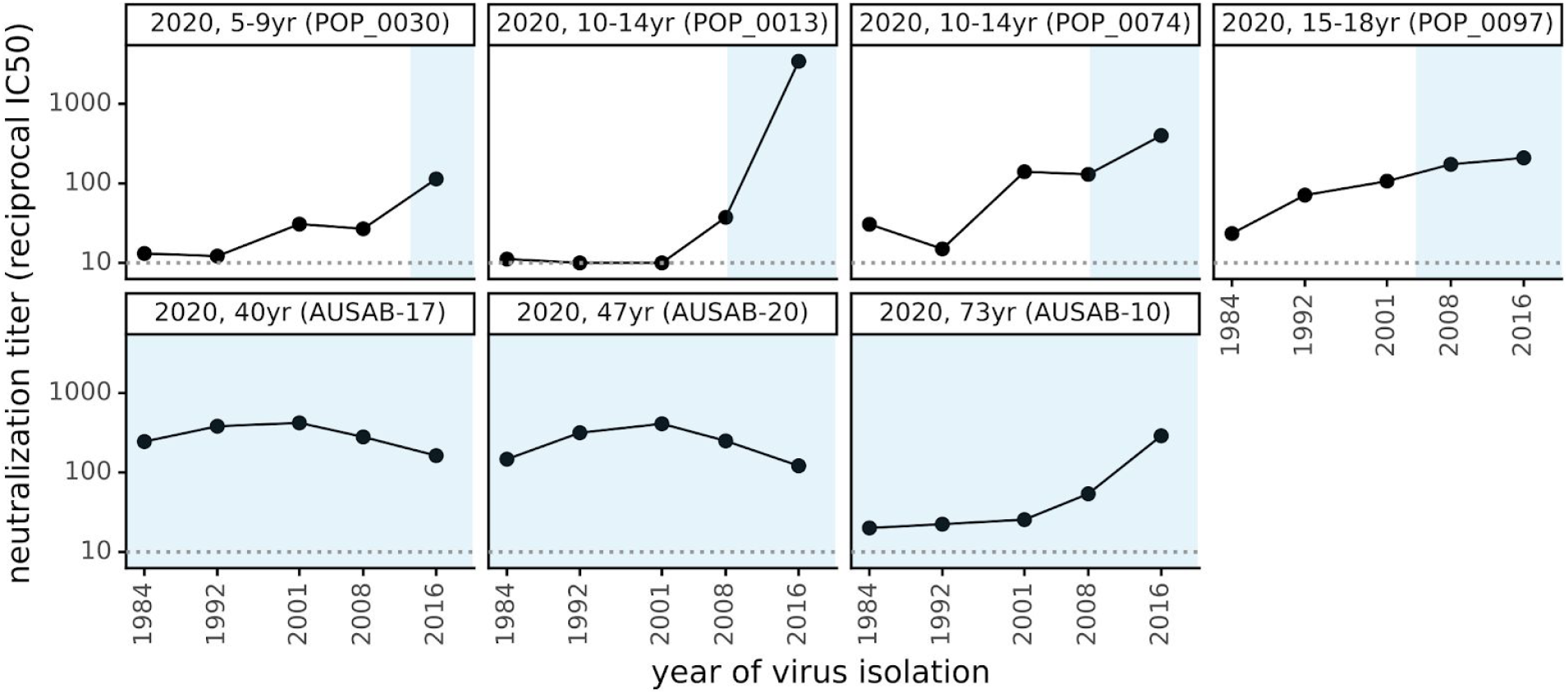
Neutralizing titers of sera collected in 2020 are higher against historical viruses that circulated during an individual’s lifetime than viruses isolated before the individual was born. As in Figure 2A,B, each plot facet is a different serum with the title giving the individual’s age and black points indicating the titer against spikes from viruses isolated in the indicated year. Blue shading represents the portion of the plotted timeframe during which the individual was alive: for adults this is the entire timeframe, but for children the left edge of the blue shading indicates the birth year.

### Much of the antigenic evolution is due to mutations in the spike’s receptor-binding domain (RBD)

We next sought to identify the region(s) within the 229E spike where mutations drive antigenic drift. Coronavirus spikes consist of two subunits, S1 and S2, and it is well known that S2 is relatively conserved whereas S1 is more rapidly evolving (Liò and Goldman, 2004; Tortorici and Veesler, 2019). The S1 subunit itself consists of several domains, and we were inspired by several excellent papers by Rini and colleagues to pursue the hypothesis that 229E’s antigenic drift might be driven by amino-acid substitutions within the three loops in the S1 RBD that bind the receptor (Li et al., 2019; Wong et al., 2017).

We first calculated the protein sequence variability at each residue across an alignment of 229E spikes isolated between 1984 and 2019 (Figure 4A). As expected, most sequence variability was in the S1 subunit, with particularly high variability in the three receptor-binding loops within the RBD (Figure 4A). However, there was also substantial variability within some portions of the N-terminal domain (NTD) as well as other parts of the S1 subunit.

**Figure 4.**
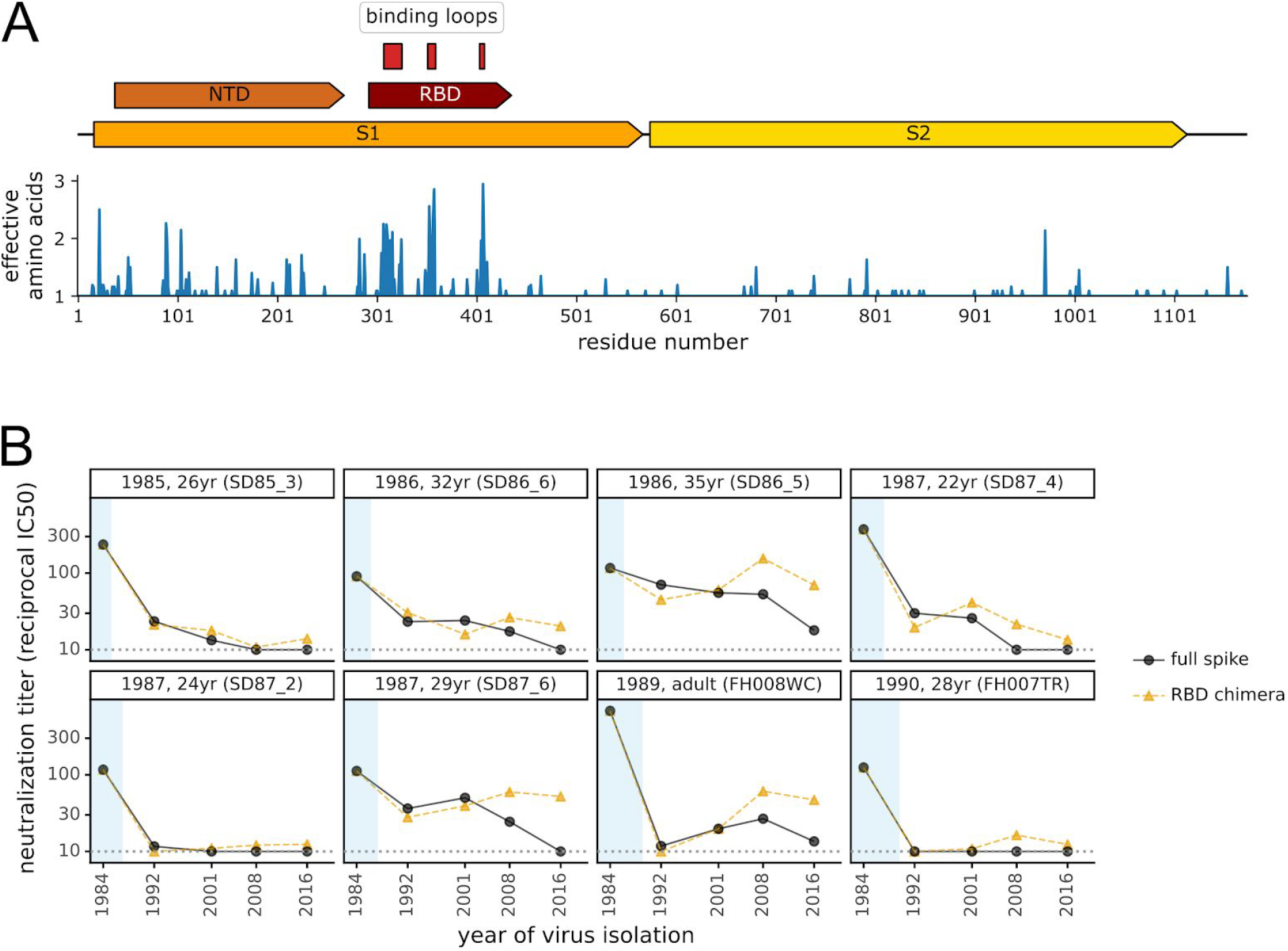
Antigenic evolution is primarily due to changes in the spike’s receptor binding domain (RBD). (A) At top is a schematic of the 229E spike. Within the S1 subunit, the schematic indicates the N-terminal domain (NTD, also known as S1-A) and the RBD (also known as S1-B). The three loops in the RBD that bind the virus’s APN receptor are indicated (Li et al., 2019). Below the schematic is a plot of sequence variability across the alignment of 229E spikes in Figure 1A. Variability is quantified as the effective number of amino acids at a site (Echave and Wilke, 2017), with a value of one indicating complete conservation and larger values indicating more sequence variability. (B) Neutralizing titers of sera collected between 1985 and 1990 against either the full spike of “future” viruses or chimeras consisting of the 1984 spike containing the RBD from “future” viruses. The plot format and the black circles (full spike) are repeated from Figure 2A with the addition of the orange triangles showing the titers against the chimeric spikes.

To experimentally test the extent that mutations in the RBD explained antigenic evolution, we created chimeras consisting of the 1984 spike with the RBD replaced by that of each of the four subsequent spikes. All these RBD-chimeric spikes supported efficient entry by pseudotyped viral particles (Figure S3). We performed neutralization assays using the chimeric spikes against the sera from 1985–1990 that potently neutralized the 1984 spike (Figure 4B). For all sera with neutralizing activity that was rapidly eroded by antigenic evolution, the drops in titer against more recent spikes were paralleled by drops in titer against the RBD-chimeric spikes (e.g., 26 and 24-year olds in Figure 4B). However, this trend did not hold for some sera that were more resistant to antigenic evolution: for instance, serum from the 29-year old did not neutralize the 2016 spike, but neutralized the chimera with the 2016 RBD (Figure 4B). Overall, these results suggest that when the neutralizing activity of potent human sera is rapidly eroded by viral evolution this is often due to mutations within the RBD, but that antigenic evolution also occurs elsewhere in the spike. In this respect, it is worth noting that while the neutralizing activity of SARS-CoV-2 immunity elicited by infection primarily targets the RBD (Piccoli et al., 2020), mutations to the NTD also reduce neutralization by some antibodies and sera (Chi et al., 2020; Kemp et al., 2020; Liu et al., 2020a; McCarthy et al., 2020; Weisblum et al., 2020; Zhou et al., 2019)—and some regions of the NTD undergo significant sequence evolution in 229E (Figure 4A).

## Discussion

We have experimentally demonstrated that the spike of a human coronavirus evolves antigenically with sufficient speed to escape neutralization by many polyclonal human sera within one to two decades. This finding suggests that one reason that humans are repeatedly re-infected with seasonal coronaviruses may be that evolution of the viral spike erodes the immunity elicited by prior infections.

How does the rate of antigenic evolution of 229E compare to that of influenza virus? Remarkably, we could find no studies that measured how quickly influenza evolution erodes neutralization by human sera. However, numerous studies examine influenza antigenic evolution using hemagglutination-inhibition (HAI) assays with sera from ferrets infected with single viral strains (Smith et al., 2004). The rate at which 229E escapes neutralization by human sera is several fold slower than the rate at which influenza A/H3N2 escapes HAI by ferret sera, but comparable to the rate of such escape by influenza B (Bedford et al., 2014; Neher et al., 2016). However, the sera of ferrets infected by a single influenza virus strain tend to recognize fewer viral strains than sera from humans who have been repeatedly infected with many strains (Fonville et al., 2014, 2016; Li et al., 2013). Therefore, ferret sera HAI may overestimate how quickly influenza evolution erodes neutralization by actual human sera. For this reason, further work is needed to enable head-to-head comparisons of antigenic evolution across these viruses.

The rapid antigenic evolution of the 229E spike might seem puzzling given that coronaviruses have a lower mutation rate than other RNA viruses (Denison et al., 2011; Sanjuán et al., 2010). But the rate of phenotypic evolution is not equal to mutation rate; evolution also depends on the effects of mutations and how selection acts on them. These other factors explain why influenza undergoes rapid antigenic evolution while measles does not, despite having a similar mutation rate. Specifically, mutations to influenza hemagglutinin are often well tolerated (Thyagarajan and Bloom, 2014), and single hemagglutinin mutations can have huge effects on escaping polyclonal sera (Lee et al., 2019). In contrast, measles surface proteins are less tolerant of mutations (Fulton et al., 2015), and the single mutations that are tolerated never more than modestly affect measles neutralization by polyclonal sera (Muñoz-Alía et al., 2020). In these respects, coronaviruses unfortunately seem more similar to influenza than measles. The neutralizing antibody response to SARS-CoV-2 is often focused on just a small portion of spike (Barnes et al., 2020; Liu et al., 2020b; Piccoli et al., 2020; Weisblum et al., 2020), and key receptor-binding loops are mutationally tolerant in the spikes of both 229E (Li et al., 2019; Wong et al., 2017) and SARS-CoV-2 (Starr et al., 2020a). Therefore, even though mutations to coronaviruses occur at a lower rate, they are acted on by selection in a fashion more similar to influenza than measles (Kistler and Bedford, 2020).

A striking aspect of our results is the extreme person-to-person variation in how rapidly neutralizing immunity is eroded by the evolution of coronavirus 229E. Some sera that potently neutralize contemporaneous virus have no detectable activity against viral strains isolated 8+ years later. But other sera maintain neutralizing activity against strains isolated over two decades later. This finding is reminiscent of how mutations to influenza virus can have vastly different effects on neutralization by sera from different individuals (Lee et al., 2019). Identifying what factors determine how rapidly an individual’s coronavirus immunity is eroded by viral mutations is an important area for future work, as it would obviously be desirable for SARS-CoV-2 vaccines to elicit immunity that is relatively robust to viral evolution.

The biggest question is what our work implies about possible antigenic evolution by SARS-CoV-2. While it is impossible to know if SARS-CoV-2 will evolve similarly to 229E, it is ominous that mutations affecting neutralization by antibodies or sera are already present at low frequencies among circulating SARS-CoV-2 (Greaney et al., 2020; Kemp, 2020; Liu et al., 2020b; McCarthy et al., 2020; Starr et al., 2020b; Thomson, 2020; Weisblum et al., 2020) even though most of the human population is still naive and so presumably exerting little immune pressure on the virus. But two facts provide hope even in light of our observation that human coronaviruses evolve to escape neutralizing immunity. First, the level of immunity required to prevent severe COVID-19 may be low (Polack et al., 2020), perhaps because the slower course of disease provides more time for a recall immune response than for “quicker” viruses such as influenza. An optimistic interpretation is that disease might often be mild even if viral antigenic evolution allows re-infections. Second, many leading SARS-CoV-2 vaccines use new technologies such as mRNA-based delivery (Krammer, 2020) that should make it easy to update the vaccine if there is antigenic evolution in spike. For this reason, we suggest that SARS-CoV-2 evolution should be monitored for antigenic mutations that might make it advisable to periodically update vaccines.

## Methods

### Data and computer code availability

The data and computer code for all phylogenetic analyses and experiments are on GitHub at https://github.com/jbloomlab/CoV_229E_antigenic_drift. Relevant parts of this GitHub repository are called out in the Methods below; in addition the repository itself includes a README that aids in navigation. All paper figures can be generated using the computer code and data in this repository.

### Phylogenetic analysis of 229E spikes

To assemble a set of 229E spikes, we downloaded all accessions for “Human coronavirus 229E (taxid:1137)” available from NCBI Virus as of July-13-2020. These accessions are listed at https://github.com/jbloomlab/CoV_229E_antigenic_drift/blob/master/data/NCBI_Virus_229E_accessions.csv. The NCBI information for some sequence accessions (particularly older sequences) were missing metadata that was available in publications describing the sequences (Chibo and Birch, 2006; Shirato et al., 2012). For these accessions, we manually extracted the relevant metadata from the publications (see https://github.com/jbloomlab/CoV_229E_antigenic_drift/blob/master/data/extra_229E_accessions_metadata.yaml). We also identified a few sequences that were clear outliers on the date-to-tip regression in the analyses described below, and so are probably mis-annotations; these accessions were excluded (see https://github.com/jbloomlab/CoV_229E_antigenic_drift/blob/master/data/accessions_to_include_exclude_annotate.yaml).

We parsed full-length human-isolate spikes encoding unique proteins from this sequence set (see https://github.com/jbloomlab/CoV_229E_antigenic_drift/blob/master/results/get_parse_spikes.md), used mafft (Katoh and Standley, 2013) to align the protein sequences, and used a custom Python script (https://github.com/jbloomlab/CoV_229E_antigenic_drift/blob/master/prot_to_codon_alignment.py) to build a codon alignment from the protein alignment (File S1). We used GARD (Kosakovsky Pond et al., 2006; Spielman et al., 2019) to screen for recombination (see tanglegram in Figure S2 and code at https://github.com/jbloomlab/CoV_229E_antigenic_drift/blob/master/results/gard_tanglegram.md).

The phylogenetic tree topology was inferred using IQ-TREE (Minh et al., 2020) using a codon-substitution model (Muse and Gaut, 1994) with a transition-transversion ratio and F3X4 empirical codon frequencies. We then used TreeTime (Sagulenko et al., 2018) to root the tree (Figure S2) and also re-scale the branch lengths for the time tree in Figure 1A. The tree images were rendered using ETE 3 (Huerta-Cepas et al., 2016).

To compute the variability at each site in spike (Figure 4A), we used the same alignment as for the phylogenetic analysis. We then computed the amino-acid variability at each site as the effective number of amino acids, which is the exponential of the Shannon entropy (Echave and Wilke, 2017). The domains of spike were annotated using the definitions in (Li et al., 2019), which are provided at https://github.com/jbloomlab/CoV_229E_antigenic_drift/blob/master/data/AAK32191_hand_annotated.gp.

### Plasmids encoding 229E spikes and RBD chimeras

The protein sequences of the spikes used in the experiments are in File S2. We deleted the last 19 amino acids of the spike’s C-terminus (in the cytoplasmic tail) as this modification has been reported (Crawford et al., 2020a; Kawase et al., 2009; Rogers et al., 2020) and validated in our hands (Figure S3) to improve titers of virions pseudotyped with spike. For the 1984, 1992, 2001, 2008, and 2016 spikes, the protein sequence matches the Genbank sequences for these viral strains (Figure 1A and File S1) except for the tail deletion. For the RBD chimeras, we annotated domains of spike as in (Li et al., 2019); see https://github.com/jbloomlab/CoV_229E_antigenic_drift/blob/master/data/AAK32191_hand_annotated.gp. We then designed the RBD-chimera proteins by replacing the RBD of the 1984 spike with the RBD of each of the 1992, 2001, 2008, and 2016 spikes.

We designed human-codon-optimized gene sequences encoding each of these spike proteins using the tool provided by Integrated DNA Technologies, had the genes synthesized commercially, and cloned them into a CMV-driven expression plasmid. Genbank sequences of the resulting plasmids are at https://github.com/jbloomlab/CoV_229E_antigenic_drift/tree/master/exptl_data/plasmid_maps. The names of the plasmids are listed below (note how the names include “delta19” or “d19” to indicate the C-terminal deletion as well as the year for that viral strain and whether it is a chimera; note also that we created a plasmid for the 2016 spike that did not have the C-terminal deletion for the experiments in Figure S3 that validated the benefits of the deletion):

- HDM-229E-Spike-d19-1984
- HDM-229E-Spike-d19-1992
- HDM-229E-Spike-d19-2001
- HDM-229E-Spike-d19-2008
- HDM-229E-Spike-Seattle2016
- HDM-229E-Spike-delta19-Seattle2016
- HDM-229E-Spike-d19-1984-1992RBD
- HDM-229E-Spike-d19-1984-2001RBD
- HDM-229E-Spike-d19-1984-2008RBD
- HDM-229E-Spike-d19-1984-2016RBD

### Generation and titering of 229E spike-pseudotyped lentiviral particles encoding luciferase and ZsGreen

We generated spike-pseudotyped lentiviral particles using the same approach that we have recently described for SARS-CoV-2 (Crawford et al., 2020a, 2020b). This approach involves creating pseudotyped lentiviral particles by transfecting cells with a plasmid expressing spike, a plasmid expressing a lentiviral backbone encoding luciferase and ZsGreen, and plasmids expressing the other lentiviral proteins necessary for virion formation (Crawford et al., 2020a, 2020b). The only modifications for this study are that we used the plasmids expressing the 229E spike described above rather than plasmids expressing the SARS-CoV-2 spike, and that after producing the pseudotyped lentiviral particles we infected them into target cells engineered to be infectable by the 229E spike.

Specifically, the 229E spike binds to human aminopeptidase N (APN) to initiate viral entry (Yeager et al., 1992). To make 293T cells infectable by 229E, we therefore transiently transfected them with an APN protein expression plasmid (SinoBiological, NM_001150.2) prior to seeding the cells for infection. To further promote lentiviral infection, we simultaneously transiently transfected them with a plasmid encoding transmembrane serine protease 2 (TMPRSS2), which facilitates 229E-spike mediated viral entry by cleaving and activating the spike (Bertram et al., 2013). We used the TMPRSS2-expressing plasmid pHAGE2_EF1aInt_TMPRSS2_IRES_mCherry (Lee et al., 2018).

For titering the spike-pseudotyped particles in these cells, we used the following procedure. To mitigate any possible well-to-well differences in transfection efficiency in a 96-well plate format, we first bulk transfected a dish of 293T cells, followed by seeding the 96-well plates routinely used in neutralization assays and viral titering. Specifically, an approximately 90% confluent 10 cm dish of 293T cells was transfected with 8.5 μg APN-expressing plasmid, 1 μg of TMPRSS2-expressing plasmid, and 0.5 μg of carrier DNA (Promega, E4881) to achieve an 8.5:1 ratio of APN:TMPRSS2. We found that this ratio gave sufficient TMPRSS2 expression, while maintaining low levels of cell toxicity. Cells were transfected using the Bioland Scientific BioT transfection reagent following the manufacturer’s protocol but incubating transfection complexes for 15 minutes at room temperature instead of the recommended 5 minutes as we have anecdotally observed that this extended incubation increases transfection efficiency. After 5–6 hours, transfection supernatant was removed and the APN and TMPRSS2-transfected 293T cells were trypsinized (Fisher, MT25053CI). Cells were then seeded in clear bottom, black-walled, poly-L-lysine coated 96-well plates that were either professionally pre-coated (Greiner, 655936) or hand-coated (Greiner, 655090) with poly-L-lysine solution (Millipore Sigma, P4704) at 1.75×10e4 cells per well in 50 μL D10 growth media (DMEM with 10% heat-inactivated FBS, 2 mM L-glutamine, 100 U/mL penicillin, and 100 μg/mL streptomycin). Plates were incubated at 37°C with 5% CO_2_ for 20–24 hours, and cells were infected with serial 2-fold serial dilutions of the pseudotyped lentiviral particles. These viral dilutions were made in TC-treated “set-up” 96-well plates and then transferred to the pre-seeded 293T-ACE2-TMPRSS2 cells from the previous day.

Approximately 50–52 hours later, we quantified infection by reading the luminescence signal produced from the luciferase encoded in the lentiviral backbone. Specifically, 100 μL of media in each well was removed—while being sure to leave the cells undisturbed—leaving approximately 30 μL of media left over. Then an equal volume of Bright-Glo reagent (Promega, E2610) was added to the remaining 30 uL of media in each well and the solution was mixed up and down to ensure complete cell lysis. In order to minimize the potential for premature luciferase excitation, special care was taken to protect the assay plates from light. Mainly, assay plate preparation was performed in a biosafety hood with the lights off and plates were covered in tin foil after the addition of the luciferase reagent. The luminescence was then measured on a TECAN Infinite M1000 Pro plate reader with no attenuation and a luminescence integration time of 1 s. Figure S3 shows the titers achieved for each 229E spike variant, and also demonstrates the importance of the spike cytoplasmic tail deletion and the expression of APN and TMPRSS2 for efficient viral infection. Note that for one panel in Figure S3, we instead determined the titer by using flow cytometry to detect the fraction of cells expressing the ZsGreen also encoded in the lentiviral backbone.

### Human sera

All sera used in this study, along with relevant metadata (e.g., collection date, patient age, and the measured neutralization titer against each assayed virus) are listed in File S3, which is also available at https://github.com/jbloomlab/CoV_229E_antigenic_drift/blob/master/exptl_data/results/all_neut_titers.csv.

Most of the historical human sera from the 1980s and 1990s are identified by the prefix SD in File S3 (e.g., SD85_1). These sera were obtained from the Infectious Disease Sciences Biospecimen Repository at the Vaccine and Infectious Disease Division (VIDD) of the Fred Hutchinson Cancer Research Center in Seattle, WA, and were collected from prospective bone marrow donors with approval from the Human Subjects Institutional Review Board. A few of the historical sera are residual samples obtained from Bloodworks Northwest that were collected from adults in Seattle; these sera are identified by the prefix FH in File S3 (e.g., FH007TR). A few of the sera were collected from subjects with exact ages that were unknown, but were adults old enough to have been alive in 1984 (the isolation year of the first spike in our panel).

The modern children’s sera from 2020 are identified by the prefix POP_ in File S3 (e.g., POP_0007), and are residual sera collected at Seattle Children’s Hospital in Seattle, WA, in March of 2020 with approval from the Human Subjects Institutional Review Board. Each of these serum samples is from a unique individual who was confirmed to be seronegative for SARS-CoV-2 by an anti-RBD ELISA (Dingens et al., 2020).

The modern adult sera from 2020 are identified by the prefix AUSAB (e.g., AUSAB-01) in File S3, and are residual sera from University of Washington Lab Medicine that were collected for testing for HBsAb (for which they tested negative).

For a negative control, we used serum from a goat that had not been infected with the 229E human coronavirus; namely WHO goat serum (FR-1377) procured from the International Reagent Resource.

All sera were heat inactivated prior to use by incubation at 56°C for approximately 30 minutes.

### Neutralization assays

The 293T cells used for our neutralization assays were transfected to express APN and TMPRSS2 and seeded as for the viral titering described above. The neutralization assays were set up at 20–24 hours after seeding of the cells into 96-well plates. First, the heat-inactivated serum samples were diluted 1:10 in D10 growth media followed by 3-fold serial dilutions in TC-coated 96-well “set-up” plates, ultimately giving seven total dilutions per sample. These dilutions were done in duplicate for each serum sample. Each 229E spike-pseudotyped lentivirus was then diluted to achieve luciferase readings of approximately 200,000 RLUs per well (the exact dilution factor varied among viruses due to differences in titers; see Figure S3). An equal volume of virus was then added to each well of the virus plus sera “set-up” plates, and these plates were incubated for 1 hour at 37°C with 5% CO_2_, after which 100 uL of each virus plus sera mixture were transferred to the 293T-APN-TMPRSS2 cell plate that had been seeded the day prior. Plates were incubated at 37°C with 5% CO_2_ for approximately 50-52 hours, and then the luciferase signal was read as described above for viral titering.

Each neutralization plate contained a column of positive control wells consisting of cells plus virus incubated with D10 media but no sera, and a negative control consisting of virus but no cells (we also confirmed that using a cells-only negative control gave similar results). The fraction infectivity was computed at each serum dilution as the fraction of the signal for the positive control (averaged across the two positive control wells for each row) after subtracting the background reading for the negative control. We then fit 2-parameter Hill curves with baselines fixed to 1 and 0 using neutcurve (https://jbloomlab.github.io/neutcurve/). Note that the serum concentration reported in these curves is the concentration at which the virus was pre-incubated with the sera for 1 hour. All of the neutralization curves are plotted in Figure S4. All of the IC50s are tabulated in File S3. See https://github.com/jbloomlab/CoV_229E_antigenic_drift/tree/master/exptl_data for raw and processed plate reader data and all of the computer code used for the fitting. Note that all assays were done in duplicate, but some sera-virus pairs have additional readings as we re-ran selected sera-virus pairs to confirm that results remained consistent across different assay days. In all cases the day-to-day consistency was good, and the reported values are the mean IC50s across all assays.

## Acknowledgments

We thank David Veesler, Allison Greaney, Tyler Starr, Kathryn Kistler, and Trevor Bedford for helpful comments. This work was supported by the NIH / NIAID under grants R01AI127983 and R01AI141707 (to JDB), and F30AI149928 (to KHDC). We also thank the Vaccine and Infectious Diseases Division of the Fred Hutch for supporting the biorepository used for the 1980s and 1990s sera. JDB is an Investigator of the Howard Hughes Medical Institute.

## Supplementary Information

**Figure S1.**
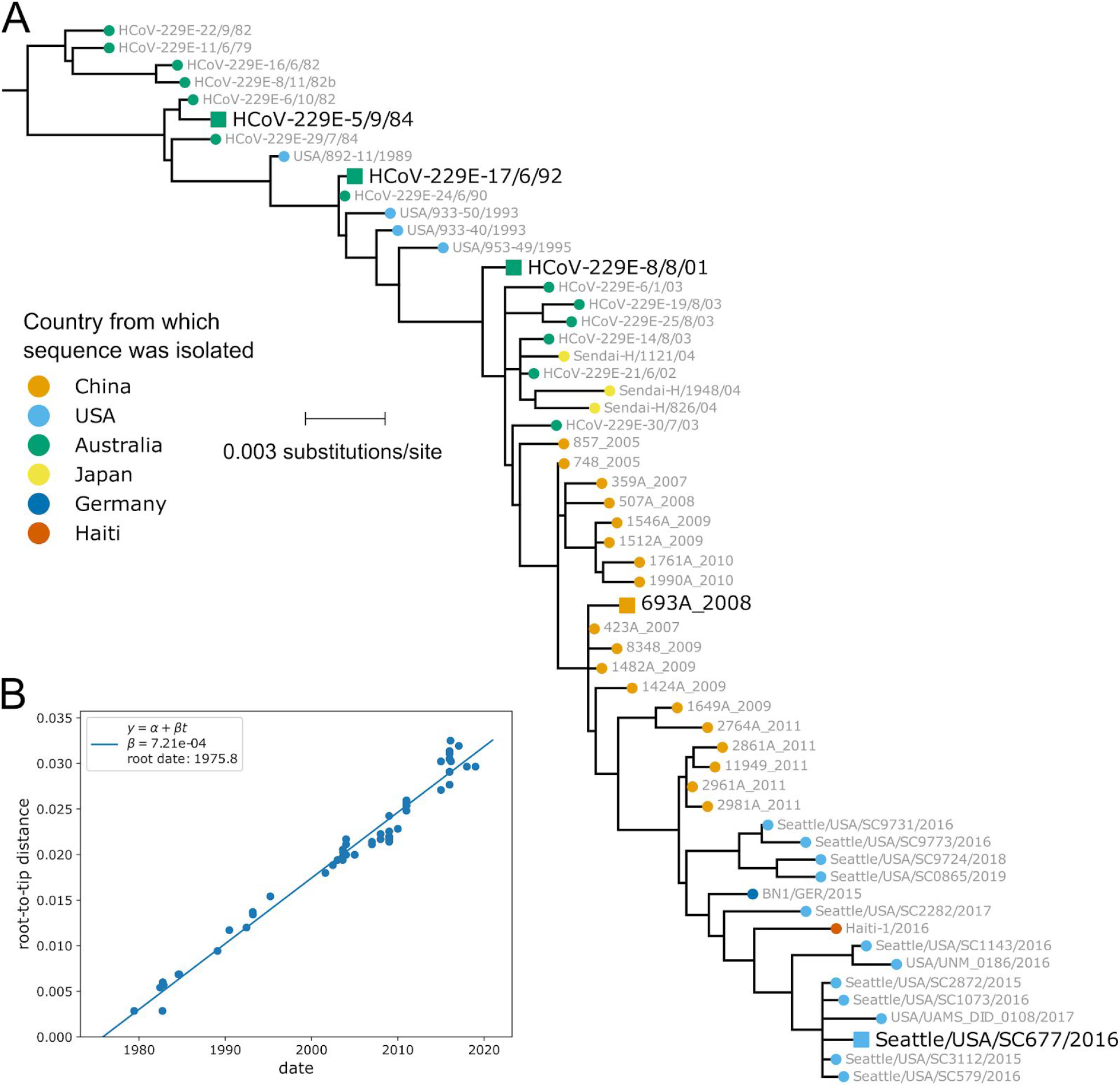
The evolution of the 229E spike is clock-like, with the number of substitutions per site proportional to time. (A) Phylogenetic tree exactly like that in Figure 1 but with branch lengths proportional to divergence (not re-scaled based on tip isolation date). (B) A plot produced by TreeTime (Sagulenko et al., 2018) showing the correlation between the distance of tip nodes from the root and sampling date. The fact that all points fall on a line indicates that the evolution is clock-like.

**Figure S2.**
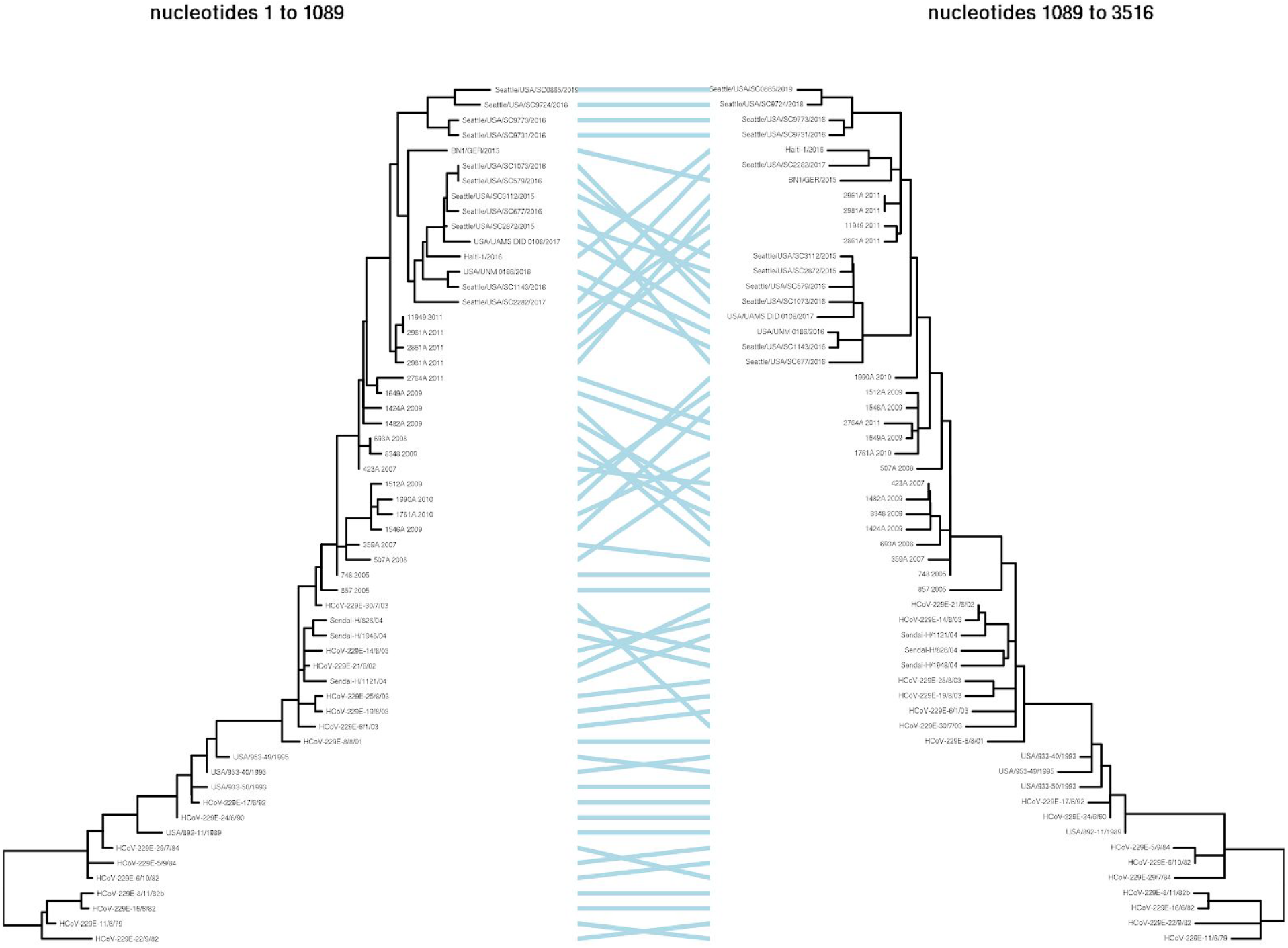
Although there is some evidence of recombination among closely related 229E spikes, this recombination does not alter the relative phylogenetic relationships among the spikes used in the experiments. Specifically, GARD (Kosakovsky Pond et al., 2006; Spielman et al., 2019) was used to analyze the same set of 229E spike sequences used in Figure 1 with a nucleotide substitution model and three gamma-distributed rate classes. The best-fitting model had a single recombination breakpoint at nucleotide 1089 that improved the AIC by 60 units. The trees for each partition were then rooted and branch-re-scaled using TreeTime (Sagulenko et al., 2018), and the resulting tanglegram was rendered using dendextend (Galili, 2015). As can be seen above, the recombination is all between closely related sequences and does not alter the relative position of the 1984, 1992, 2001, 2008, and 2016 spikes used in the experiments. See https://github.com/jbloomlab/CoV_229E_antigenic_drift/blob/master/results/gard_tanglegram.md for details of the analysis steps described above.

**Figure S3.**
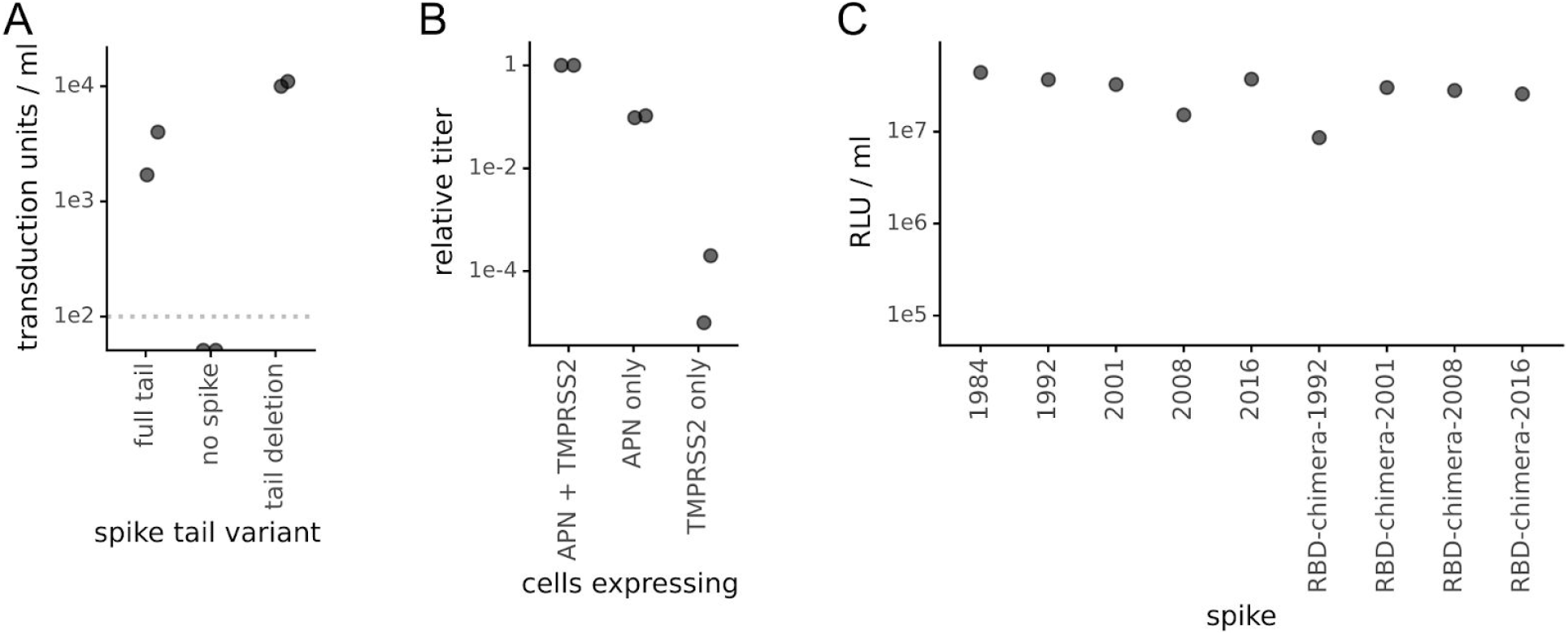
The 229E spikes with a cytoplasmic tail deletion pseudotype lentiviral particles that efficiently infect 293T cells expressing the spike’s receptor aminopeptidase N (APN) and the activating protease TMPRSS2. (A) Titer in transduction units per ml as determined using flow cytometry of lentiviral particles pseudotyped with the full-length 2016 spike or that spike with a deletion of the last 19 residues in spike (the end of the cytoplasmic tail) on 293T cells transfected with a plasmid expressing APN. The dotted gray line is the limit of detection, and the titers in the absence of spike were below this line (undetectable). (B) Efficient entry by the pseudotyped virions depends on expression of APN and to a lesser extent TMPRSS2. Virions pseudotyped with the 2016 spike with the C-terminal deletion were infected into 293T cells transfected with plasmids expressing one or both of APN and TMPRSS2, and titers were determined by luciferase luminescence. Titers are normalized to one. (C) All of the 229E spikes and chimeras used in this study mediated efficient viral entry. Lentiviral particles were pseudotyped with the indicated spike (in all cases with the C-terminal deletion) and titers were determined using luciferase luminescence on 293T cells transfected with plasmids expressing APN and TMPRSS2.

**Figure S4.**
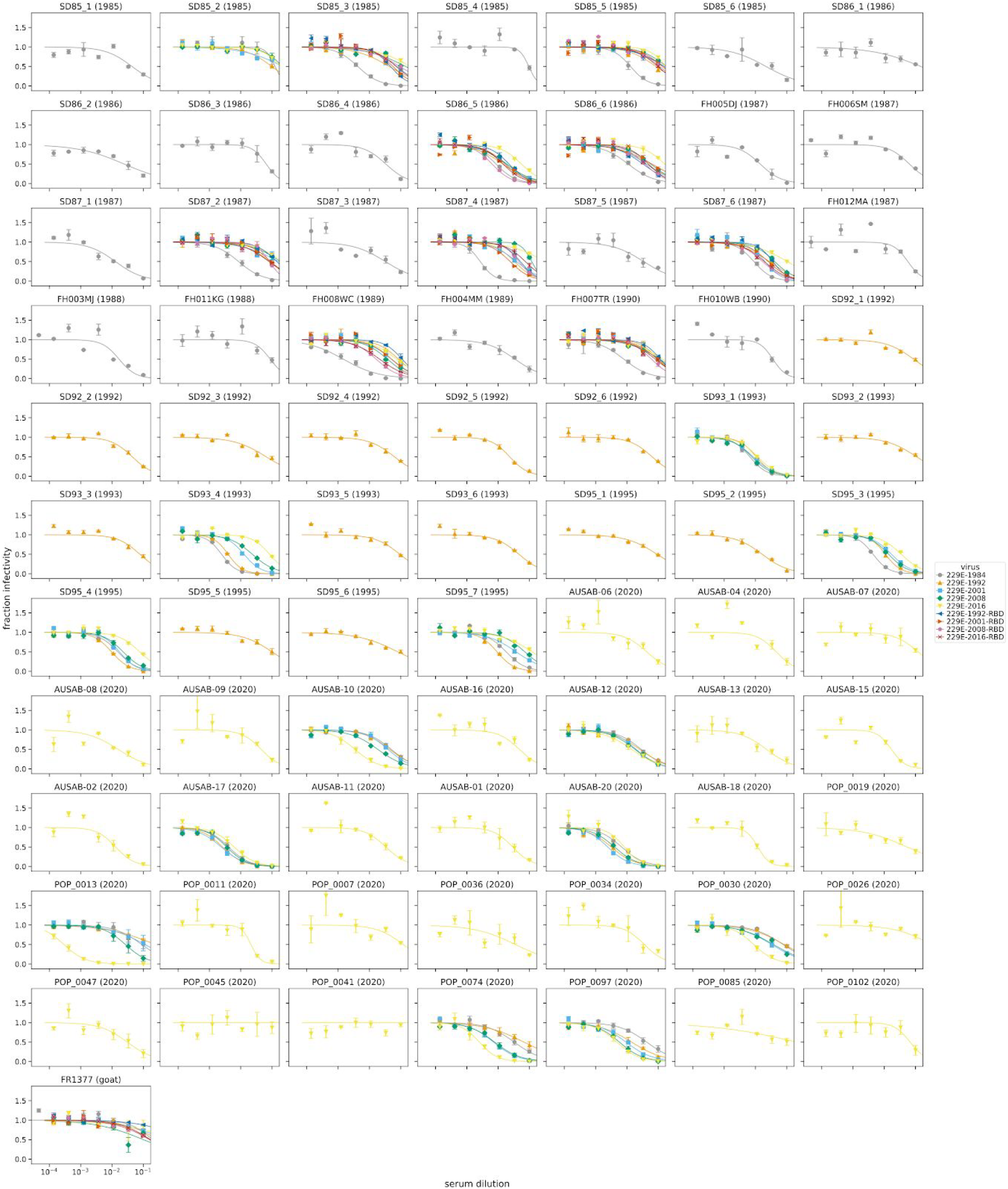
Neutralization curves for all assays. Each facet is a serum, with titles indicating the year the serum was collected. Each point is the fraction infectivity at that serum concentration averaged across at least two replicates (error bars are standard error), with colors indicating the virus. The fits are 2-parameter Hill curves with baselines fixed to 1 and 0, and were fit using neutcurve (https://jbloomlab.github.io/neutcurve/). IC50s are in File S3. The curves are also at https://github.com/jbloomlab/CoV_229E_antigenic_drift/blob/master/exptl_data/results/all_neut_by_sera.pdf

**Figure S5.**
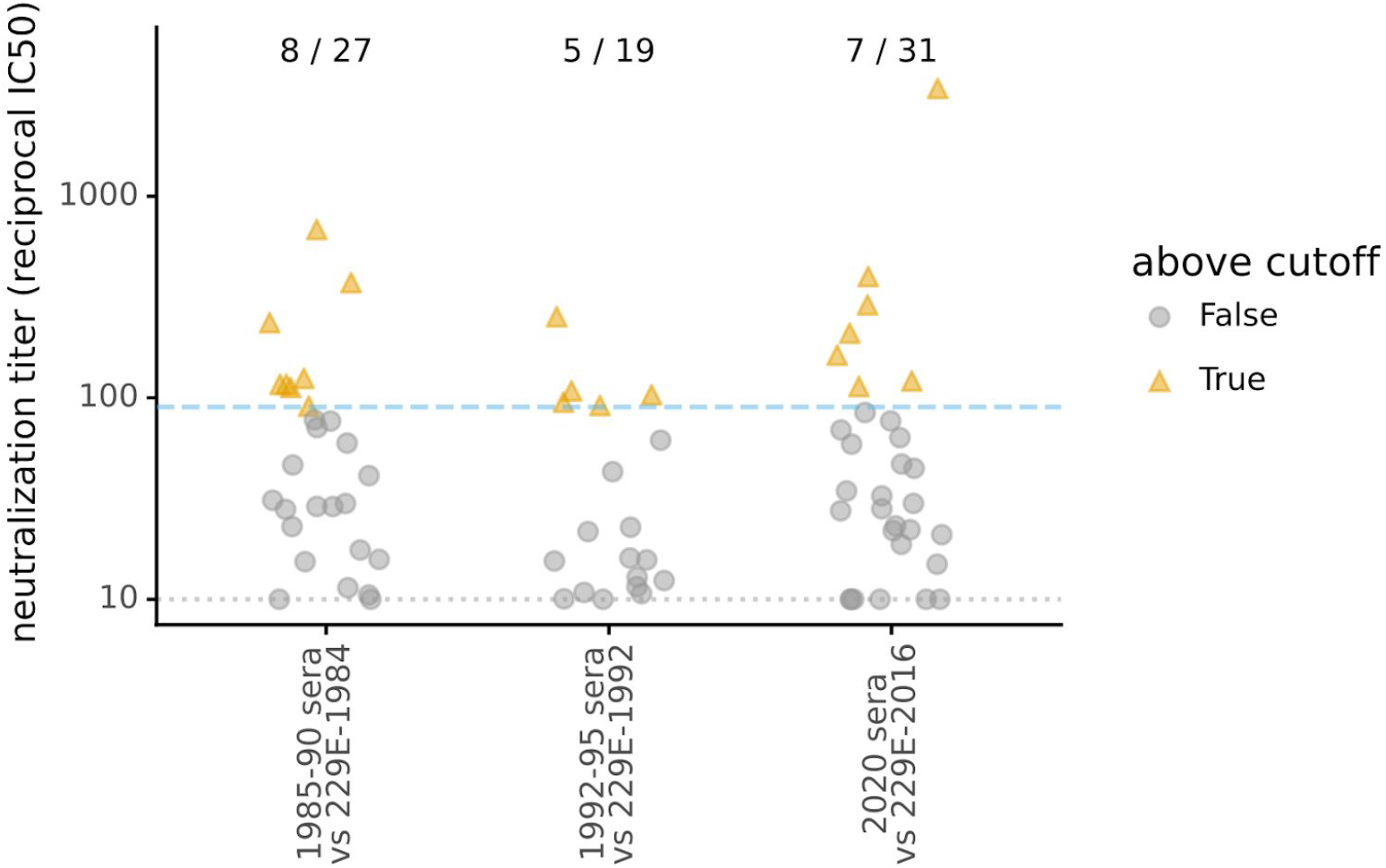
Initial screening of sera to identify samples with neutralizing titers of at least 1:90 that were then used for the rest of the studies described in the paper. Each sera was tested against the most-recent virus isolated prior to the serum collection date: in other words, sera collected between 1985–1990 was tested against the 1984 spike, sera collected between 1992–1995 was tested against the 1992 spike, and sera collected in 2020 was tested against the 2016 spike. Each point shows the neutralizing titer for a different serum (see Figure S4 for full neutralization curves). Sera above the cutoff of 1:90 (blue dashed line) was then used for further studies against the full panel of viruses (e.g., Figures 2, 3, and 4). The numbers at the top of the plot indicate the number of sera above the cutoff out of the total sera tested in each timeframe. The dotted horizontal line at the bottom of the plot is the lower limit of detection of the neutralization assay. Quantitative neutralization titers for each sera are in File S3.

---

**File S1.** Codon-level alignment of the 229E spike sequences. This FASTA alignment is at https://github.com/jbloomlab/CoV_229E_antigenic_drift/blob/master/results/spikes_aligned_codon.fasta

**File S2.** A ZIP of GenPept files giving the protein sequences of the spikes used in the experiments. There are nine sequences: the five spikes from the 1984, 1992, 2001, 2008, and 2016 viruses (named by strain as shown in Figure 1A), and the four chimeras that consist of the 1984 spike with the RBD of each of the other strains. Each GenPept file annotates key domains in the spike. Note that the C-terminal 19 amino acids are deleted off each spike. These files are at https://github.com/jbloomlab/CoV_229E_antigenic_drift/tree/master/results/seqs_for_expts

**File S3.** A CSV file giving the neutralization titer, collection date, and subject age at time of collection date for each serum sample analyzed in this study. This file is at https://github.com/jbloomlab/CoV_229E_antigenic_drift/blob/master/exptl_data/results/all_neut_titers.csv

